# Structural Insights into Single-Stranded DNA Recognition and Modified Substrate Tolerance in an Engineered Terminal Deoxynucleotidyl Transferase

**DOI:** 10.64898/2025.12.25.696546

**Authors:** Lixiang Yang, Hanyi Liao, Qiyue Wang, Weiwei Li, Chang Xu, Jianxun Qi, Huijun Zhang, Lifeng Fu, Meng Yang

**Affiliations:** MGI Tech, Shenzhen 518083, China; Laboratory of Pathogen Microbiology and Immunology, Institute of Microbiology, Chinese Academy of Sciences, Beijing 100101, China; Faculty of Health Sciences, University of Macau, Macau, SAR, 999078, China; Medical School, University of Chinese Academy of Sciences, Beijing 101408, China

## Abstract

Terminal deoxynucleotidyl transferase (TdT) is the cornerstone enzyme for de novo enzymatic DNA synthesis (EDS), promising to overcome the length and sustainability limitations of traditional phosphoramidite chemistry. We previously identified a highly active TdT ortholog from Zonotrichia albicollis (ZaTdT) and engineered a variant (R335L/K337G) capable of efficiently incorporating 3’-aminooxy (3’-ONH_2_) reversible terminators. However, the atomic-level mechanism by which these engineered mutations alter substrate specificity has remained elusive. Here, we present the high-resolution (2.20 Å) crystal structure of the engineered ZaTdT in complex with a single-stranded DNA primer. Structural analysis reveals a conserved catalytic core anchored by a hydrophobic platform (Phe110/Phe257/Trp308) that stabilizes the primer. Crucially, comparative modeling with the homologous murine TdT ternary complex demonstrates that the engineered mutations (corresponding to L190/G192 in the crystal structure) disrupt a rigid salt-bridge network at the active site entrance. This electrostatic remodeling not only reduces local positive charge but also expands the catalytic pocket depth from 3.8 Å to 5.8 Å. This specific spatial expansion provides the structural rationale for the accommodation of bulky 3’-blocking groups, validating our rational design strategy and paving the way for next-generation long-read DNA synthesis.

## 1. Introduction

The ability to synthesize long, high-fidelity DNA sequences is a fundamental driver of the bioeconomy, underpinning advancements in synthetic biology, metabolic engineering, and DNA data storage [1–3]. For decades, the nucleoside phosphoramidite method has served as the gold standard; however, it is constrained by a practical length limit (∼200 nt), the accumulation of toxic byproducts, and a reliance on anhydrous conditions [4, 5]. Enzymatic DNA synthesis (EDS) offers a transformative alternative, operating under mild aqueous conditions with the potential for synthesis speeds and lengths that far exceed chemical methods [6].

Terminal deoxynucleotidyl transferase (TdT), a unique template-independent polymerase of the Pol X family, is the primary engine for EDS [7]. TdT naturally catalyzes the random addition of nucleotides to single-stranded DNA (ssDNA) during V(D)J recombination [8]. To repurpose TdT for precise, sequence-defined synthesis, the reaction must be controlled using 3’-O-blocked reversible terminators (RTs) that prevent uncontrolled polymerization [9, 10]. While various blocking groups such as 3’-O-azidomethyl and 3’-O-aminooxy (3’-ONH_2_) have been developed [11], wild-type TdTs—typically of murine or bovine origin—exhibit poor catalytic efficiency towards these unnatural substrates due to steric hindrance within the active site [12].

In our previous work, we screened TdT orthologs from diverse vertebrates and identified Zonotrichia albicollis TdT (ZaTdT) as a superior candidate with intrinsic activity significantly higher than mammalian homologs [13]. Through rational design, we introduced specific mutations (R335L/K337G) that conferred the ability to incorporate 3’-ONH_2_-dNTPs, enabling the stepwise synthesis of defined oligonucleotides. Building on this, we recently applied advanced computational strategies, including ProteinMPNN and PROSS, to address the enzyme’s thermal instability. We successfully generated a hyper-stable variant (M7-8) that retains high activity at elevated temperatures, facilitating the synthesis of difficult sequences prone to secondary structures [14].

Despite these biochemical and functional breakthroughs, the structural mechanism governing the engineered ZaTdT’s substrate tolerance has been inferred solely from homology modeling. The lack of experimental structural data hinders further optimization of the enzyme for bulkier modifications or faster kinetics. In this study, we bridge this gap by solving the crystal structure of the engineered ZaTdT complexed with ssDNA. Our findings reveal how specific mutations reshape the electrostatic and steric landscape of the catalytic pocket, providing a definitive structural explanation for the enzyme’s enhanced performance in EDS cycles.

## 2. Materials and Methods

### 2.1 Protein Expression and Purification

The coding sequence of ZaTdT (146-513) with two mutations, R335L and K337G, was cloned into a pSKB2 containing a C-terminal His_6_ tag. The proteins were expressed in E. coli BL21 (DE3) cells. Protein expression was induced with 0.2 mM IPTG at an OD_600_ of 0.6–0.8, followed by incubation at 16 °C for 16–18 h. Cells were then harvested by centrifugation (8,000 rpm, 10 min, 4 °C) and resuspended in Buffer A (20 mM Tris-HCl pH 8.0, 500 mM NaCl, 25 mM imidazole, 0.1 mM TCEP). The suspension was lysed by sonication on ice. Cell debris was removed by centrifugation at 12,000 rpm for 30 min at 4 °C. The supernatant was applied to a Ni-NTA column pre-equilibrated with Buffer A. After loading, bound proteins were eluted using Buffer A supplemented with 250 mM imidazole. To remove the C-terminal His_6_ tag, the proteins were incubated with HRV 3C protease overnight at 4 °C. Tag-free proteins were purified by SEC on a Superdex 75 10/300 GL column equilibrated with SEC buffer (20 mM Tris-HCl pH 8.0, 150 mM NaCl). The purified proteins were concentrated to 10 mg/mL and stored at -80 °C.

### 2.2 Crystallization and Data Collection

Crystallization was achieved using the sitting-drop vapor diffusion method at 18 °C. Purified ZaTdT (10 mg/mL) was mixed with a single-stranded DNA primer (5’-dTdTdTdTdT-3’) at a molar ratio of 1:1.5 and incubated at 4 °C for 30 minutes. Crystals appeared after two weeks in drops containing 0.1 M BIS-TRIS (pH 6.5) and 25% (v/v) polyethylene glycol 300. Crystals were flash-frozen in liquid nitrogen. Diffraction data were collected at the Shanghai Synchrotron Radiation Facility (SSRF).

### 2.3 Structure Determination

Data were processed and scaled using HKL2000. The structure was solved by molecular replacement using the murine TdT structure (PDB ID: 4I27) as a search model. Refinement was carried out using Phenix and Coot. The final model was validated using MolProbity.

## 3. Results

### 3.1 Crystallographic Analysis of the ZaTdT-ssDNA Complex

We successfully crystallized the engineered ZaTdT variant (R335L/K337G) in complex with ssDNA. The crystal diffracted to a resolution of 2.20 Å and belonged to the monoclinic space group P 2_1_. The final structure was refined to R_work and R_free values of 23.61% and 28.83%, respectively, with excellent stereochemical quality (96.23% residues in favored Ramachandran regions). Detailed data collection and refinement statistics are summarized in Table 1.

**Table 1.** Crystallographic data collection and refinement statistics.

ZATdT mut + ssDNA
|  |  |
| --- | --- |
| **Data collection** |  |
| Space group | P 21 |
| Cell dimensions |  |
| *a*, *b*, *c* (Å) | 52.265, 135.853, 67.984 |
| *α*, *β*, *γ* (°) | 90.000, 90.226, 90.000 |
| Wavelength (Å) | 0.97853 |
| Resolution (Å) | 50.00-2.20 (2.28-2.20) |
| Unique reflections | 47434 |
| *R*<sub>merge</sub> | 0.098 (1.284) |
| *I* / *σI* | 19.06 (21.68) |
| Completeness (%) | 84.68 |
| Redundancy | 6.5 (6.5) |
| **Refinement** |  |
| Resolution (Å) | 34.22-2.20 |
| No. of reflections | 40824 |
| *R*<sub>work</sub> / *R*<sub>free</sub> | 0.2361/0.2883 |
| No. of atoms |  |
| Protein | 676 |
| Ligand/ion | 12 |
| Water | - |
| B-factor |  |
| Protein | 38.53 |
| Ligand/ion | 46.33 |
| Water | - |
| R.m.s.deviation |  |
| Bond length (Å) | 0.0047 |
| Bond angles (°) | 0.74 |
| Ramachandran analysis |  |
| Most favored (%) | 96.23 |
| Allowed (%) | 3.77 |
| Disallowed (%) | 0.00 |

\*values in parentheses are for highest-resolution shell.

### 3.2 Overall Architecture and Primer Stabilization

The solved structure reveals that ZaTdT adopts the classic “right-hand” polymerase fold (Fig. 1b). Sequence alignment (Fig. 1a) and structural superposition with murine TdT (PDB: 4I27) demonstrate a high conservation of the catalytic core (RMSD < 1.0 Å). The electron density map allows for unambiguous tracing of the ssDNA substrate (Fig. 1e). Crucially, we identified a robust interaction network stabilizing the primer strand (Fig. 1f). The phosphate backbone is anchored by salt bridges with Arg290 and Lys115, along with hydrogen bonds involving Thr116 and Gly113. Distinctively, aromatic residues Phe110, Phe257 and Trp308 form a “hydrophobic platform” that sandwiches the terminal bases, ensuring the 3’-OH is correctly oriented for catalysis. This explains the high affinity of ZaTdT for ssDNA initiators observed in our previous kinetic studies [13].

**Figure 1.**
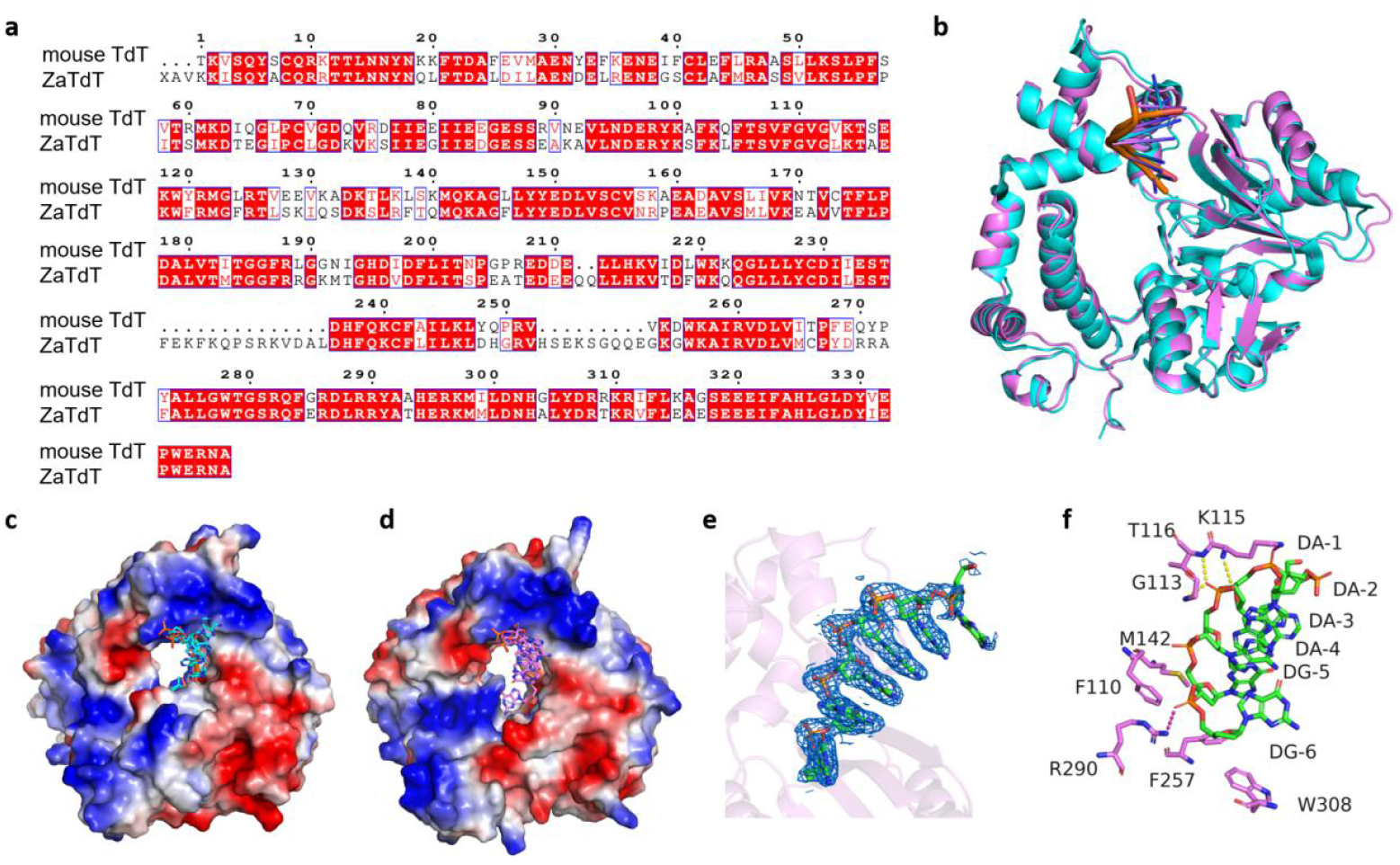
Structural and Sequence Analysis of ZaTdT. (a) Sequence alignment of the truncated ZaTdT construct used for crystallization and the murine TdT (mTdT). Conserved residues are highlighted in red boxes. (b) Structural superposition of the solved ZaTdT structure (violet) and the reference mTdT structure (cyan, PDB ID: 4I27), illustrating the highly conserved overall fold of the catalytic core. (c) Electrostatic potential surface representations of mTdT (left) and ZaTdT (right). Blue and red surfaces indicate positive and negative potentials, respectively. The single-stranded DNA (ssDNA) substrates are depicted as sticks in cyan (mTdT model) and violet (ZaTdT structure), occupying the positively charged DNA-binding groove. (e) Electron density map (blue mesh) of the ssDNA substrate bound within the ZaTdT active site, showing well-defined conformation.(f) Detailed view of the molecular interactions between ZaTdT residues and the ssDNA substrate. The phosphate backbone is stabilized through salt bridges with Arg290 and Lys115, and hydrogen bonds involving Thr116, Lys115, and Gly113 (yellow dashed lines). The aromatic residues Phe110, Trp308 and Phe257 form a hydrophobic platform providing structural support for the RNA/DNA terminal bases.

### 3.3 Structural Mechanism of 3’-ONH_**2**_ Accommodation

The primary barrier to enzymatic DNA synthesis (EDS) using reversible terminators is the steric exclusion of 3’-blocked nucleotides by the polymerase active site. As we did not obtain a crystal structure of the wild-type ZaTdT, we utilized the high-resolution ternary complex of the homologous murine TdT (mTdT, PDB ID: 4I27) as a structural reference for the catalytically competent, “closed” state. By comparing the mTdT structure with our engineered ZaTdT variant, we identified three key structural alterations that facilitate the acceptance of 3’-ONH_2_-dNTPs:

1. Electrostatic Remodeling: Electrostatic potential surface analysis highlights a distinct difference between the two enzymes. The active site entrance of mTdT is characterized by a dense cluster of positive charges, which creates a constrained electrostatic environment (Figure 2a). In contrast, the engineered ZaTdT variant—specifically at the mutation sites L190 and G192 (corresponding to R335L/K337G in the full sequence)—displays a significantly neutralized surface potential and a visibly enlarged cavity opening (Figure 2c). This reduction in positive charge density likely lowers the electrostatic barrier for the entry and positioning of modified substrates.
2. Disruption of the Salt-Bridge Network: In the mTdT structure, the triphosphate moiety of the incoming nucleotide is rigidly locked in place by a network of salt bridges formed by residues Arg336 and Lys338 (Figure 2b). To evaluate the impact of our engineering, we modeled the position of the incoming nucleotide in the ZaTdT active site by structural superposition with 4I27. This model reveals that the mutation of these key residues to Leucine (L190) and Glycine (G192) in ZaTdT completely abolishes these salt-bridge interactions (Figure 2d). While the base orientation remains stabilized by conserved hydrogen bonds with Arg312 and Arg319, the loss of the phosphate-clamping interactions introduces necessary local flexibility, allowing the substrate to adjust to the presence of the 3’-modification.
3. Catalytic Cavity Expansion: Structural superposition reveals the most critical consequence of the mutations: a conformational shift in the backbone loop containing L190 and G192. In mTdT, this loop constricts the space near the 3’-hydroxyl group to approximately 3.8 Å. In the engineered ZaTdT, the replacement of bulky basic side chains with the smaller Leucine and Glycine residues widens this gap to 5.8 Å (Figure 2e). This additional 2.0 Å of spatial clearance provides sufficient volume to accommodate the bulky 3’-aminooxy (3’-ONH_2_) moiety without steric clashes, serving as the structural basis for the efficient incorporation of reversible terminators.

**Figure 2.**
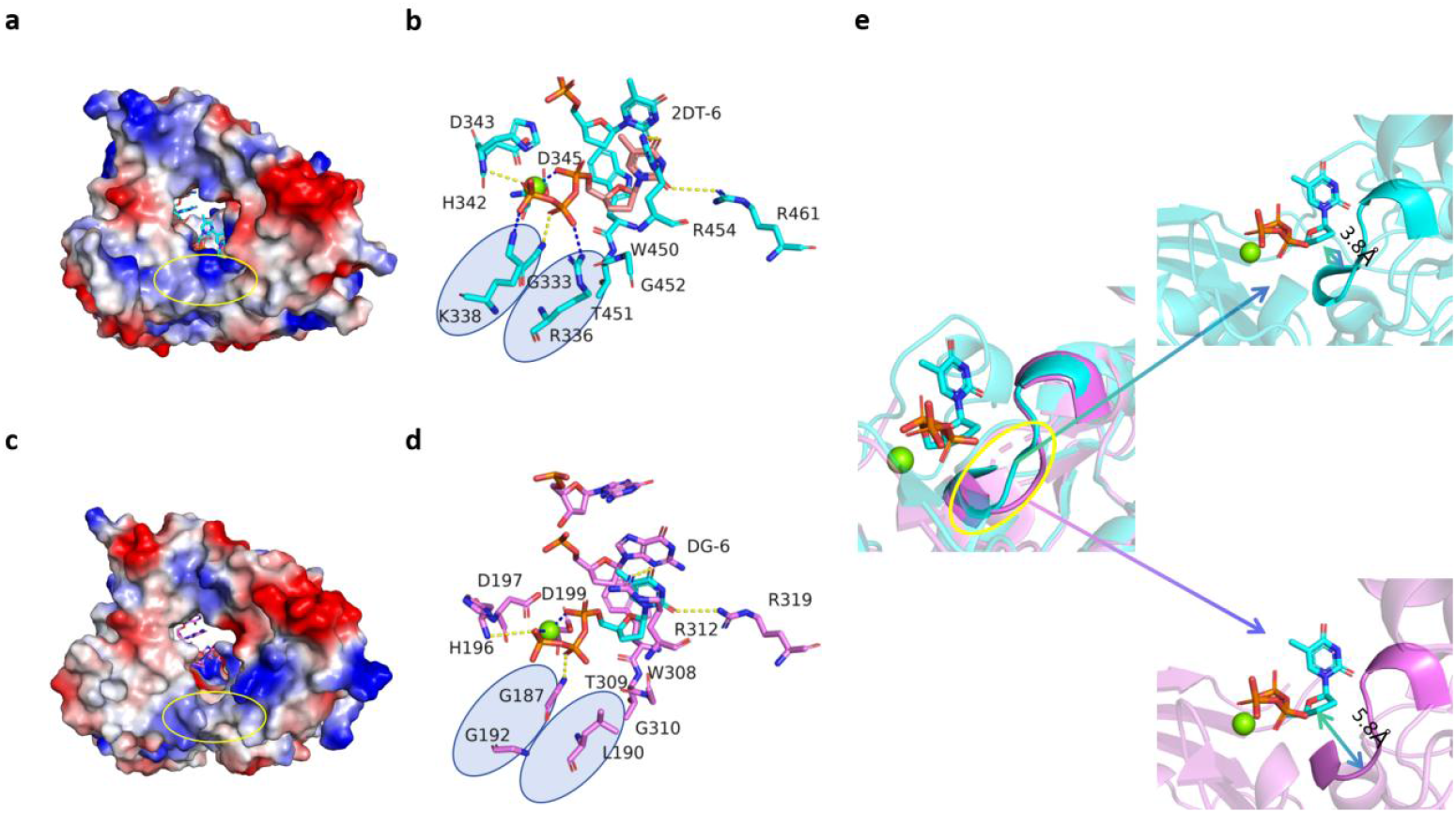
Structural mechanism of 3’-ONH_**2**_ substrate accommodation in engineered ZaTdT. (a, c) Comparison of electrostatic potential surfaces between murine TdT (mTdT, PDB: 4I27) (a) and the engineered ZaTdT structure (c). Blue and red represent positive and negative potentials, respectively. The yellow circles highlight the mutation site entrance. The mutations of positively charged residues to Leucine and Glycine (L190/G192 in the crystal structure) in ZaTdT result in a significantly reduced positive charge and an enlarged cavity compared to mTdT, increasing flexibility for substrate entry. (b, d) Detailed view of the catalytic center and substrate interactions. (b) The active site of mTdT (cyan) showing the incoming nucleotide (ddTTP) stabilized by a network of salt bridges with residues K338, R336, and others (blue ovals). (d) The active site of the engineered ZaTdT (violet). Note: The substrate shown is modeled by structural superposition with the mTdT ternary complex (PDB: 4I27) to visualize potential interactions. The mutations R190L and K192G (corresponding to R335L/K337G in the full sequence) abolish the salt bridges with the triphosphate group (blue ovals), leading to increased local flexibility. Key hydrogen bonds with the base (via R312, R319) remain conserved. (e) Structural superposition focusing on the 3’-end of the incoming nucleotide. The loop region containing the mutations undergoes a conformational shift. The distance available near the 3’-position expands from 3.8 Å in mTdT (cyan) to 5.8 Å in the engineered ZaTdT (violet). This widened spatial clearance (indicated by the arrow) is sufficient to accommodate the bulky 3’-aminooxy (3’-ONH_2_) modification, serving as the structural basis for modified substrate incorporation.

## 4. Discussion

The structural data presented here unify the findings of our previous work on ZaTdT discovery [13] and thermostability design [14]. While computational tools like ProteinMPNN and PROSS successfully stabilized the protein scaffold, the specific enzymatic activity towards modified substrates is dictated by the rational engineering of the active site.

Our structure clarifies that residues Phe110, Trp308 and Phe257 are responsible for the stable binding of the primer strand, consistent with their high conservation and structural rigidity. In contrast, the substrate specificity for the incoming nucleotide is governed by the engineered loop containing L190/G192 (mutations R335L/K337G). The transition from a “tight, charged” pocket to a “spacious, neutral” one is the key mechanism enabling EDS with reversible terminators. This aligns with the “steric gate” concept in polymerase engineering but highlights a unique mechanism involving both steric debulking and electrostatic relaxation [15, 16].

In conclusion, this study provides the atomic-level blueprint of an engineered TdT capable of synthesizing DNA with modified nucleotides. Combined with our thermostable variants, ZaTdT represents a robust platform for the realization of long-read, enzymatic DNA synthesis.

## Data availability

The atomic coordinates and structure factors for the two A Fab–ligand complexes have been deposited with the Worldwide Protein Data Bank (wwPDB). Accession codes will be provided upon release.

## Acknowledgements

This research is supported by Ministry of Science and Technology of the People’s Republic of China’s program titled ‘National Key Research and Development Program of China’ (No. 2022YFF1202200, 2022YFF1202203) and Science, Technology, Innovation Commission of Shenzhen Municipality under grant No. JSGGZD20220822095802006, the Youth Innovation Promotion Association CAS (2023096).

## Author contribution

M.Y. and L.F. conceived and designed the study. L.X.Y. and L.F. performed analysis and wrote the manuscript. L.F. contributed to the design and guidance of the experimental section. H.L., Q.W., W.L. and C.X. performed protein expression and purification and crystallization. J.Q. collected the structural data and solved the structures. M.Y. provided suggestions for the writing and revision of the manuscript. All authors read and approved the final version of the manuscript.

## References

1. Kosuri, S., & Church, G. M. (2014). Large-scale de novo DNA synthesis: technologies and applications. Nature Methods, 11(5), 499–507.

2. Ceze, L., Nivala, J., & Strauss, K. (2019). Molecular digital data storage using DNA. Nature Reviews Genetics, 20(8), 456–466.

3. Hughes, R. A., & Ellington, A. D. (2017). Synthetic DNA synthesis and assembly: putting the synthetic in synthetic biology. Cold Spring Harbor Perspectives in Biology, 9(1), a023812.

4. Caruthers, M. H. (2011). Gene synthesis machines: DNA chemistry and its uses. Science, 334(6061), 1353.

5. Roy, B., Depaix, A., Périgaud, C., & Peyrottes, S. (2016). Recent trends in nucleotide synthesis. Chemical Reviews, 116(18), 10277–10327.

6. Palluk, S., Arlow, D. H., de Rond, T., Barthel, S., Kang, J. S., Bector, R., … & Keasling, J. D. (2018). De novo DNA synthesis using polymerase-nucleotide conjugates. Nature Biotechnology, 36(7), 645–650.

7. Motea, E. A., & Berdis, A. J. (2010). Terminal deoxynucleotidyl transferase: the story of a misguided DNA polymerase. Biochimica et Biophysica Acta (BBA)-Proteins and Proteomics, 1804(5), 1151–1166.

8. Gouge, J., Rosario, S., Romain, F., Beguin, P., & Delarue, M. (2013). Structures of intermediates along the catalytic cycle of terminal deoxynucleotidyltransferase: dynamical aspects of the two-metal ion mechanism. Journal of Molecular Biology, 425(22), 4334–4352.

9. Chen, F., Dong, M., Ge, M., Zhu, L., Ren, L., Liu, G., & Mu, R. (2013). The history and advances of reversible terminators used in new generations of sequencing technology. Genomics, Proteomics & Bioinformatics, 11(1), 34–40.

10. Lee, H. H., Kalhor, R., Goela, N., Bolot, J., & Church, G. M. (2019). Terminator-free template-independent enzymatic DNA synthesis for digital information storage. Nature Communications, 10(1), 2383.

11. Hutter, D., Kim, M. J., Karalkar, N., Leal, N. A., Chen, F., Guggenheim, E., … & Benner, S. A. (2010). Labeled nucleoside triphosphates with reversibly terminating aminoalkoxyl groups. Nucleosides, Nucleotides and Nucleic Acids, 29(11), 879–895.

12. Barthel, S., Palluk, S., Hillson, N. J., Keasling, J. D., & Arlow, D. H. (2020). Enhancing terminal deoxynucleotidyl transferase activity on substrates with 3′ terminal structures for enzymatic de novo DNA synthesis. Genes, 11(1), 102.

13. Lu, X., Li, J., Li, C., Lou, Q., Peng, K., Cai, B., … & Ma, Y. (2022). Enzymatic DNA synthesis by engineering terminal deoxynucleotidyl transferase. ACS Catalysis, 12(5), 2988–2997.

14. Niu, Y., Chen, B., Zhang, H., Zheng, W., Wu, J., Yang, L., … & Yu, H. (2025). Computational Design of a Thermostable and Highly Active Terminal Deoxynucleotidyl Transferase for Synthesis of Long De Novo DNA Molecules. ACS Catalysis, 15, 7201–7216.

15. Delarue, M., Boulé, J. B., Lescar, J., Expert-Bezançon, N., Jourdan, N., Sukumar, N., … & Papanicolaou, C. (2002). Crystal structures of a template-independent DNA polymerase: murine terminal deoxynucleotidyltransferase. The EMBO Journal, 21(3), 427–439.

16. Moon, A. F., Garcia-Diaz, M., Batra, V. K., Beard, W. A., Bebenek, K., Kunkel, T. A., … & Pedersen, L. C. (2007). The X family portrait: structural insights into biological functions of X family polymerases. DNA Repair, 6(12), 1709–1725.

17. Ashley, J., Schaap-Johansen, A. L., Mohammadniaei, M., Naseri, M., Marcatili, P., Prado, M., & Sun, Y. (2021). Terminal deoxynucleotidyl transferase-mediated formation of protein binding polynucleotides. Nucleic Acids Research, 49(2), 1065–1074.

18. Loc’h, J., Rosario, S., & Delarue, M. (2016). Structural basis for a new templated activity by terminal deoxynucleotidyl transferase: implications for V (D) J recombination. Structure, 24(9), 1452–1463.

19. Romain, F., Barbosa, I., Gouge, J., Rougeon, F., & Delarue, M. (2009). Conferring a template-dependent polymerase activity to terminal deoxynucleotidyltransferase by mutations in the Loop1 region. Nucleic Acids Research, 37(14), 4642–4656.

20. Jumper, J., Evans, R., Pritzel, A., Green, T., Figurnov, M., Ronneberger, O., … & Hassabis, D. (2021). Highly accurate protein structure prediction with AlphaFold. Nature, 596(7873), 583–589.

21. Mathews, A. S., Yang, H., & Montemagno, C. (2016). Photo-cleavable nucleotides for primer free enzyme mediated DNA synthesis. Organic & Biomolecular Chemistry, 14(35), 8278–8288.

22. Jensen, M. A., & Davis, R. W. (2018). Template-independent enzymatic oligonucleotide synthesis (TiEOS): its history, prospects, and challenges. Biochemistry, 57(12), 1821–1832.

